# metaAPA: a tool for integration of PolyA site predictions from single-cell and spatial transcriptomics

**DOI:** 10.1101/2025.09.29.679268

**Authors:** Qian Zhao, Magnus Rattray

## Abstract

**Motivation:** Single-cell 3’-tagging sequencing, such as that provided by 10x Genomics, can be utilized to study alternative polyadenylation (APA). APA can affect RNA function, stability, and subcellular localization, thereby influencing development and disease processes. Currently, computational tools based on various algorithms, such as Sierra, polyApipe, and SCAPE, have been developed to infer polyA site positions from scRNA-seq data. However, these methods exhibit significant differences in the number of predicted sites and positional inconsistencies in the sites identified for the same gene, leading to divergent conclusions when analyzing the same data with different tools.

**Results:** We designed two strategies to integrate the outputs of alternative APA tools, enabling users to select appropriate polyA site sets based on their specific needs. Our method can be used to extract high confidence sites, supported by all methods, as well as putative sites supported by a subset of methods. We find that methods with high sensitivity for detecting APA sites can be usefully augmented by methods with higher positional accuracy but lower sensitivity. We show that our method obtains the expected number of high-confidence sites and that these sites exhibit the expected biological sequence characteristics.

**Availability and Implementation:** https://github.com/ManchesterBioinference/metaAPA

## Introduction

Alternative polyadenylation (APA) refers to the process by which transcripts originating from the same gene can add polyA tails at different positions through the regulation of various factors during transcription, generating transcripts with different 3’ ends. This process can increase transcript diversity, regulate RNA stability, affect subcellular localization, and is widely present in eukaryotes [1]. Popular single-cell and spatial transcriptomics methods based on 3’-tagging sequencing, such as those provided by 10x Genomics, are well-suited to identify polyA sites, thereby facilitating the study of the biological functions of APA events at the single-cell level.

Alignment-based methods (mapping reads by Cell Ranger, Space Ranger or STARsolo instead of Kallisto or Salmon) for identifying polyA sites in scRNA-seq can be mainly divided into four types [2]. The first type identifies peaks by fitting the distribution of coverage, represented by scAPA [3], Sierra [4], and SCAPTURE [5]. The second type infers the position of the site by estimating the distance from the read to the polyA site, represented by MAAPER [6] and SCAPE [7]. The third type identifies change points within a given region to segment peaks and obtain site information without a distribution prior, represented by scDarPars [8]. The fourth type of method identifies polyA sites by obtaining reads with a polyA tail through the soft-clip mechanism in sequence alignment, represented by polyApipe (the algorithm is unpublished, but there is an application published in Cell [9]), SCINPAS [10], and scTail [11]. Additionally, some methods attempt to combine the different mechanisms mentioned above to enhance their performance, such as scAPAtrap [12] (combining the third and fourth type) and Infernape [13] (combining the first and second type).

Through benchmarking, we have observed that different algorithms exhibit varying performance [2]. For instance, SCAPE, a method of the second type, can identify low-coverage sites but is susceptible to the effects of alternative splicing [2, 7]. Sierra, a method of the first type, accounts for alternative splicing but requires sufficient reads to form peaks for fitting [2, 4]. The soft-clip-based polyApipe method may lose true signals or identify false positives due to sequencing depth and read length, although it provides excellent results when sufficient biological sequence data are available [2, 9]. Integrating sites identified by methods based on different algorithms could potentially enhance the strengths and mitigate the weaknesses of each, providing a more comprehensive APA landscape. However, due to algorithmic differences, the number of sites identified for the same sample can vary significantly, and the inferred positions of these sites often exhibit discrepancies, making direct comparison or integration challenging.

Given that different tools perform well under different scenarios, it can be beneficial to consider evidence from multiple approaches. Some methods attempt to combine two different algorithms by using one primary method with another as a filter to enhance site identification precision. However, this filtering approach is typically not as optimized as a standalone tool. Furthermore, methods that integrate three or more principles have not yet been developed. We have therefore designed a flexible integration framework that can combine outputs from different methods based on two distinct integration strategies (Fig. 1A and B). We show that this ensemble approach leads to improved precision for high-confidence sites while also providing greater sensitivity to uncover putative sites that have less uniform support across methods. Another advantage is that methods with higher positional accuracy but lower coverage can usefully augment less precise methods with higher coverage.

**Fig. 1.**
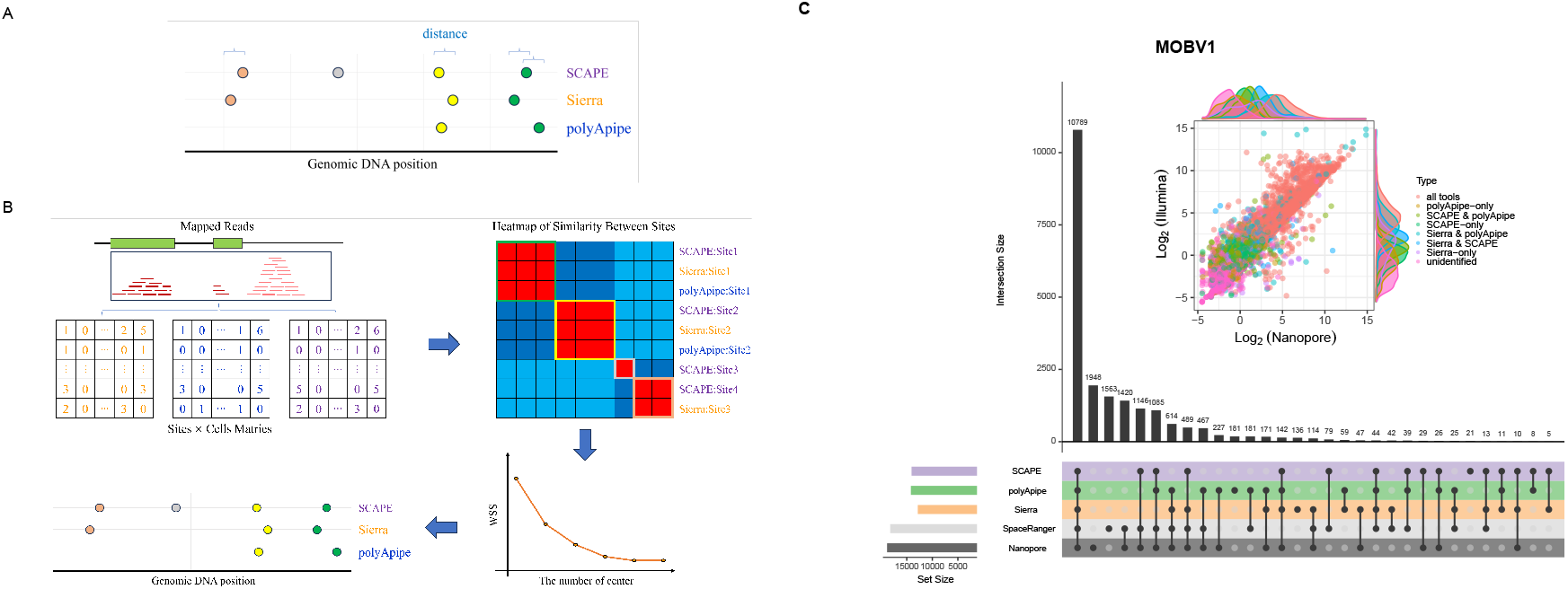
Integrative strategies for polyA site identification. (A) Position-based strategy. Calculate the distance between genomic coordinates of sites from different sources, grouping sites with distances below a specified threshold into clusters, with each cluster representing a true signal. (B) Similarity-based strategy. Perform unsupervised clustering of sites from different sources based on expression similarity. After optimizing the number of clustering centers, the resulting clusters represent the integrated results. (C) The upset plot shows the overlap in the number of genes identified by the three representative methods, Space Ranger, and Nanopore in the same dataset MOBV1. The internal scatter plot and marginal density plot illustrate the expression distribution of different categories of genes (classified by the methods that successfully identified them) among the high-confidence genes detected by both Space Ranger and Nanopore. Specifically, “all tools” refers to genes identified by all three APA analysis tools, while “unidentified” indicates genes that were not detected by any of the tools. Gene expression levels are presented as log-transformed, normalized expression values.

## Method

### Datasets

Public scRNA-seq (10x Chromium), spatial transcriptomics (10x Visium), and ONT data were collected from NCBI (see Table 1 for details). The E18, MOBV1, CBS1 and CBS2 datasets are hybrid single-cell or spatial transcriptomics sequencing datasets, meaning they used the same sequencing library for both long-read and short-read sequencing. MOBV1, CBS1, and CBS2 are three spatial transcriptomics datasets that utilized the original BAM files as input (pre-launch version 4509.7.5 of Space Ranger). E18 is a single-cell sequencing dataset, for which we remapped the FASTQ files using a different version of Cell Ranger (version 7.0.1) than the original study (version 2.0.0) to obtain the BAM file.

**Table 1.** Datasets used in the paper.

| Sample name | Type | Cells/Spots | Depth | Read length | GEO | Ref |
| --- | --- | --- | --- | --- | --- | --- |
| Mouse Brain E18 (E18) | 10X Chromium | 1159 | 71M | 132 | GSE130708 | [14] |
| Mouse Brain E18 (E18) | Nanopore | 1159 | 57M | - | GSE130708 | [14] |
| Mouse Olfactory Bulb (MOBV1) | 10X Visium | 918 | 254M | 91 | GSE153859 | [15] |
| Mouse Olfactory Bulb (MOBV1) | Nanopore | 918 | 25M | - | GSE153859 | [15] |
| Mouse Brain (CBS1) | 10X Visium | 2560 | 217M | 91 | GSE153859 | [15] |
| Mouse Brain (CBS1) | Nanopore | 2560 | 38M | - | GSE153859 | [15] |
| Mouse Brain (CBS2) | 10X Visium | 2500 | 210M | 91 | GSE153859 | [15] |
| Mouse Brain (CBS2) | Nanopore | 2500 | 91M | - | GSE153859 | [15] |

### APA identification tools

We selected three representative methods, Sierra (version), polyApipe (version), and SCAPE (version), so as to include results from a diverse set of algorithmic approaches. Following the tutorials for each method, we configured the environments and ran these methods on our data. The results include both the predicted polyA site locations and the corresponding single-cell expression matrix. We annotated the genomic coordinates of the predicted sites to retrieve their associated gene information. Subsequently, we reformatted the outputs of the three methods to generate annotated site-by-cell/spot matrices. We filtered these matrices using the filtered barcodes generated by Cell Ranger or Space Ranger, and then removed sites that were not detected in any valid cells.

To validate the scalability of the integration framework, we selected SCAPE-APA [16], a recently released improved version of SCAPE, as an additional APA detection tool. The same processing steps were applied accordingly.

### Position-based integration

To integrate sites derived from different methods based on genomic coordinates, we used the cluster module from BEDtools (version 2.31.1). First, we converted the sites identified by the three methods into a 6-column BED format (chromosome, start, end, strand, name, score), then merged and sorted them. The “name” field includes information such as the APA tool, gene name, and peak ID, while the “score” field contains the total UMI count for the site. Next, we used the BEDtools cluster command to integrate nearby sites, ensuring strand consistency (parameter *−s*) and adjusting the distance parameter (parameter *−d*). This command aggregates sites within a given threshold distance into a cluster. To verify the impact of different distances on the number of clusters, we extended the distance threshold from 0 to 3000 to evaluate the relationship between distance and performance.

### Expression similarity-based integration

Given the absence of tools that integrate sites based on similarity of expression pattern, we designed a workflow to perform this task. First, we extracted sites identified by the three methods for each gene and then calculated the expression similarity of all sites between each pair of methods. The resulting similarity matrix was clustered, with the clusters representing the integrated results. Since there are various methods for calculating similarity, choosing clustering methods, and optimizing clustering, we tested different combinations to determine the most suitable for our task. We calculated the Euclidean distance, Spearman correlation coefficient, cosine distance and Jaccard index for assessing similarity. For clustering methods, we used k-means clustering, k-medoids clustering, HDBSCAN clustering and hierarchical clustering (ward linkage).

For k-means and k-medoids, we determined the optimal number of clusters using either the within-cluster sum of squares (WSS) or the average silhouette width (ASW). For hierarchical clustering, we constrained the number of sites within each cluster to not exceed the number of APA detection tools used (*n* = 3). In cases where multiple clustering outcomes were equivalent, we prioritized clusters composed of sites identified by different tools over those formed solely by a single method.

### Evaluating the performance of site integration

Due to the lack of sufficiently accurate ground truth in polyA site integration research, we defined the number of high-confidence sites as a metric to evaluate integration effectiveness. Our hypothesis is that if a cluster contains three sites, each identified by different algorithms, the site is more likely to represent a true signal, making the cluster a high-confidence cluster and the sites within it high-confidence sites. Conversely, if a cluster contains only one site, it is considered method-specific.

Theoretically, every gene captured in sequencing should have produced at least one RNA molecule with a polyA tail (or polyA-tail-like structure), as this is required for oligo-dT-based sequencing. The goal of APA analysis tools is to identify these RNA molecules with distinct polyadenylation sites, determining whether a gene utilizes a single or multiple polyA sites (i.e., whether it gives rise to structurally distinct RNA isoforms). By annotating the genomic coordinates of the polyA sites identified by these tools, we can retrieve the corresponding gene names. Similar to the concept of high-confidence sites, if a gene is identified by all three methods, it is more likely to be truly expressed, which means it has produced at least one detectable RNA molecule with a polyA tail. Therefore, the intersection of genes identified by all three methods represents a set in which each gene should have at least one polyA site. Consequently, the lower bound for the number of polyA sites in this context can be estimated as the number of overlapping genes identified by all three methods, multiplied by three (*N*_high-confidence sites_ ≥ *N*_APA method_ ×*N*_overlapping genes_) where *N*_APA method_ = 3 refers to the three methods used. If more methods are used then this value needs to be adjusted accordingly.

In addition to the number of high-confidence sites, we also compared the overlap of high-confidence sites identified by different strategies using the Jaccard index. Consistent performance across methods should yield higher values. Furthermore, to demonstrate that high-confidence sites are more likely to represent true signals compared to method-specific sites, we extracted sequences 100bp upstream and downstream of the sites and compared their nucleotide composition for typical sequence characteristics, such as the polyadenylation signal (PAS; typically AAUAAA or its variants, located around 21 nucleotides upstream of the cleavage site), the cleavage site itself (commonly CA), the CSTF recognition motif (a G/U-rich region downstream of the cleavage site), and the CF I recognition motif (UGUA, typically located 40 nt upstream of the cleavage site) [1].

## Results

### Different APA methods identify sites from different gene sets

To illustrate the necessity of data integration, we first assess differences in site identification among the different tools. Due to the positional shifts in the sites inferred by these tools, it is challenging to directly compare the overlap of the sites or to determine which sites are missing. To address this issue, we chose to compare at the gene level rather than the site level. The identified sites, once annotated, can be assigned to the corresponding genes, and the presence or absence of genes is generally more reliable and stable.

As shown in Fig. 1C, the genes identified by the three tools exhibit varying degrees of absence. Since the input file for the three APA identification tools is BAM file generated by Space Ranger (SR), if a gene can be identified by SR but not by any APA identification tools, it indicates that the current tools are missing at least as many sites as the number of genes. Additionally, although ONT has a lower sequencing depth compared to Illumina, it offers significant advantages in transcript alignment and identification. Since ONT data are derived from the same sequencing library, if a gene is supported by both SR and ONT, it is more likely to be present in the current sample. Similarly, if some genes are identified by both SR and ONT but not by one or more APA tools, we consider that these tools have likely missed at least this number of sites. For example, we found that 614 genes were identified by all methods except SCAPE, 489 genes were identified by all methods except polyApipe, and 1085 genes were identified by all methods except Sierra (Fig. 1C). These results reflect that there are indeed differences in the identification capabilities of the current methods, with each method having genes that are uniquely identified as well as genes that are uniquely missed.

We further examined the gene composition of those supported by both SR and ONT data to investigate whether there were any patterns in gene detection or omission (Fig. 1C). We observed that genes undetected by any method were predominantly lowly expressed, whereas genes with relatively high expression levels were consistently identified by all tools. Compared to SCAPE and polyApipe, Sierra appeared to be more sensitive to expression levels, showing limited detection capability for genes with lower abundance.

### Clustering sites by genomic distance

After observing differences in the sites identified by different tools, we proposed two strategies to integrate the sites (Fig. 1 A and B). The first approach involves clustering based on genomic position (position-based strategy). Specifically, when sites overlap or the distance between sites is less than a threshold, these sites are considered to originate from the same polyA site. This strategy is straightforward and simple.

To implement the position-based strategy we used the cluster module of BEDtools to integrate these sites. Since BEDtools performs clustering solely based on genomic proximity without considering gene annotations, there is a risk of incorrect clustering, particularly when using larger distance thresholds. For example, if site *a* belongs to gene *A* and site *b* to gene *B*, and genes *A* and *B* are partially overlapping or adjacent, the distance between sites *a* and *b* may fall below the threshold, resulting in their erroneous assignment to the same cluster. To mitigate such cases, we re-group the BEDtools output based on gene annotations, ensuring that each cluster contains only sites originating from the same gene.

In Fig. 2A we have categorized clusters based on the number of sites they contained, including: single-site clusters, two-site clusters, three-site clusters, medium clusters (4-6), and large clusters (11-20). When the distance threshold was set to zero, most sites did not overlap, resulting in only a few two- or three-site clusters, and no medium or large clusters. As the distance increased, the number of single-site clusters decreased significantly, with the rate of decline slowing after approximately 50nt. Once the threshold exceeded 200nt, the rate of increase of medium-sized clusters increases, and large clusters appeared beyond 270nt. This trend suggests that increasing the distance threshold may lead to more frequent over-clustering events, where unrelated sites are grouped together, potentially compromising the performance of the integration.

**Fig. 2.**
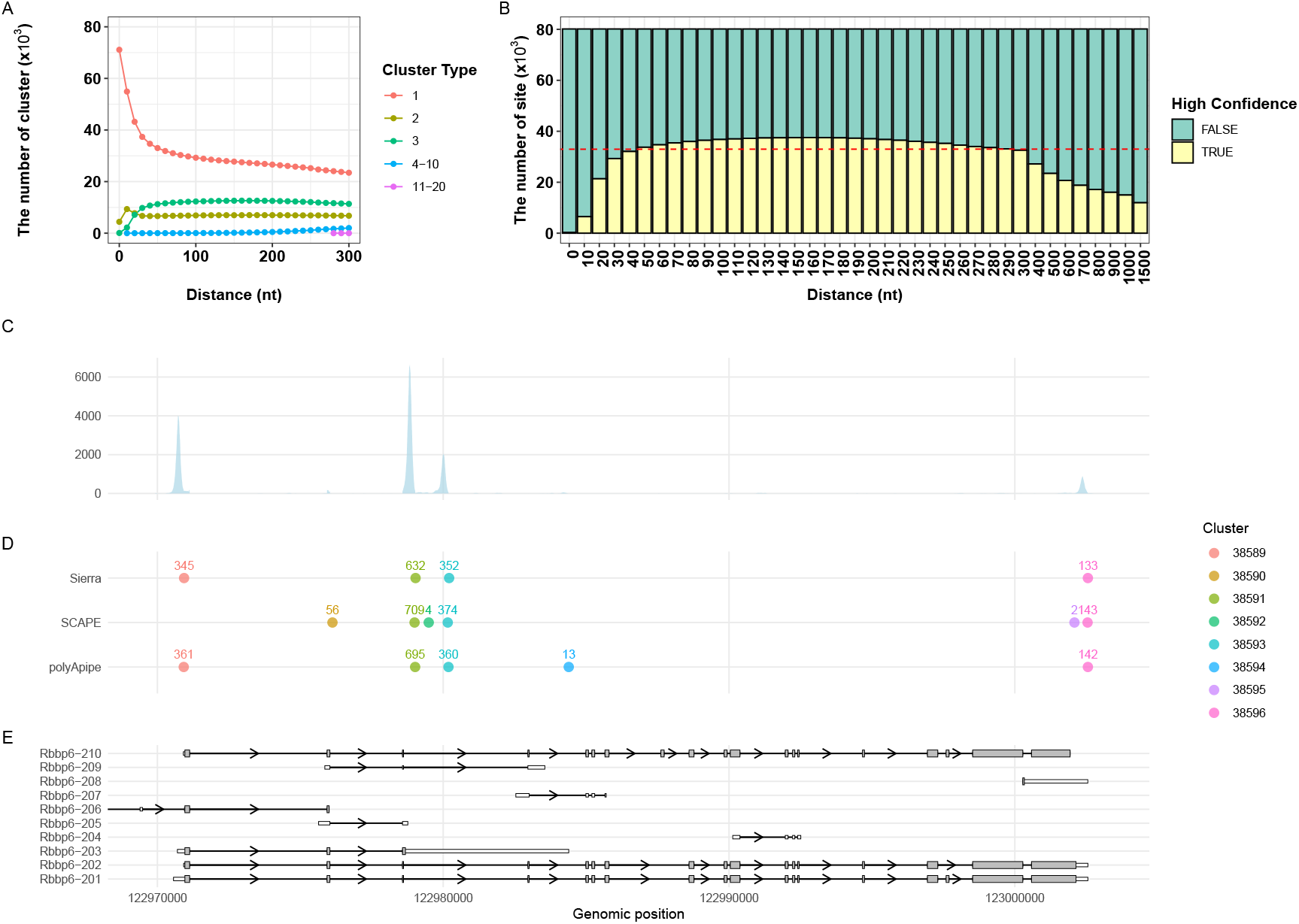
Performance of the position-based method implemented using BEDtools. A. The number of clusters in each size category against distance threshold. B. The number of high-confidence sites against distance threshold. The red dashed line indicates the high-confidence site threshold. C. Coverage plot of Rbbp6 gene. D. Clustering results for Rbbp6 gene (using distance threshold 160nt); numbers indicate number of UMIs associated with each site. E. Transcript structure of Rbbp6 gene.

To evaluate the performance of the integration and identify the optimal distance, we compared the changes in high-confidence sites with varying distances. High-confidence sites are defined as clusters containing exactly three sites, each supported by a different method. These three sites are considered high-confidence sites, and the corresponding cluster is termed a high-confidence cluster. Given that the number of genes overlapping is 10,986, the theoretically shared sites should be no less than 32,958 (3 *×* 10, 986). As shown in the Fig. 2B, the number of high-confidence sites increases with distance, exceeding 32,958 at 50nt. The growth rate slows at 50nt but continues to increase, reaching a maximum at 160nt (37,329). Subsequently, the number of high-confidence sites begins to decrease, with the rate of decrease accelerating as the distance further increases. Although the sites predicted by the three methods should be located near the same 3’ UTR, an increased distance threshold is required to achieve accurate integration due to positional shifts. This result aligns with our expectations. However, when the distance exceeds a certain range, sites from different 3’ UTRs are clustered into the same group, reducing the number of high-confidence sites (Fig. 2B).

Fig. 2C-E provides an example of the clustering results (with distance threshold 160nt) for the retinoblastoma binding protein 6 (Rbbp6) gene. This gene belongs to the ubiquitin E3 ligase family and has multiple isoforms with different functions. The position-based integration strategy identified three high-confidence clusters in the MOBV1 sample, making it a suitable case for demonstration. From the read coverage (Fig. 2C) we observe three distinct peaks that correspond to the positions of the three high-confidence clusters (clusters 37606, 37608, 37611). Their abundance decreases sequentially, consistent with the UMI results. Based on the genomic annotation of this gene, peak 1 (the highest peak) is most likely derived from Rbbp6-205 or 203, peak 2 (the second highest peak) is most likely from Rbbp6-203, and peak 3 (the third highest peak) is most likely from Rbbp6-201, 202, or 208. In the matching ONT data, we found relatively high expression of Rbbp6-203 (127 UMIs) and Rbbp6-205 (190 UMIs), as well as very low expression of Rbbp6-209 (2 UMIs), partially validating that the high-confidence sites identified by this strategy may be supported by true transcripts. The transcript corresponding to peak 3 may not have been detected by ONT due to its low expression abundance in the library.

### Clustering sites based on expression similarity

Our second strategy to integrate the sites discovered by different methods is based on expression similarity. Since the input for each method is the same BAM file, if different methods identify sites from the same signal source then these sites should exhibit high correlation across cells or spots even if there are large positional deviations. Therefore, we can achieve integration by clustering sites from different tools based on the similarity of their expression levels (where expression refers to the number of reads mapped to each site).

The workflow implementing this strategy is illustrated in Fig. 1B. Since co-expression may occur between sites originating from different genes, high correlation across genes could interfere with the clustering of sites within the same gene. To address this, we extracted expression matrices for sites originating from the same gene across all tools and merged them. The resulting expression matrices were then used to calculate similarity between these sites. The similarity matrices served as input for unsupervised clustering, and the resulting clusters were considered the final integrated output.

Using Rbbp6 as an example, Fig. 3A shows the cosine similarity between sites and sites with higher similarity are clustered together. Subsequently, we applied k-means clustering to the similarity matrix of these sites and used WSS (within-cluster sum of square) to evaluate the performance differences in the number of clusters, thereby determining the optimal number of centers (Fig. 3B). Here, based on WSS, we identified the optimal number of centers as 6. The clustering results for this gene were similar to those obtained by the first strategy, particularly in identifying the same high-confidence sites (Fig. 3C).

**Fig. 3.**
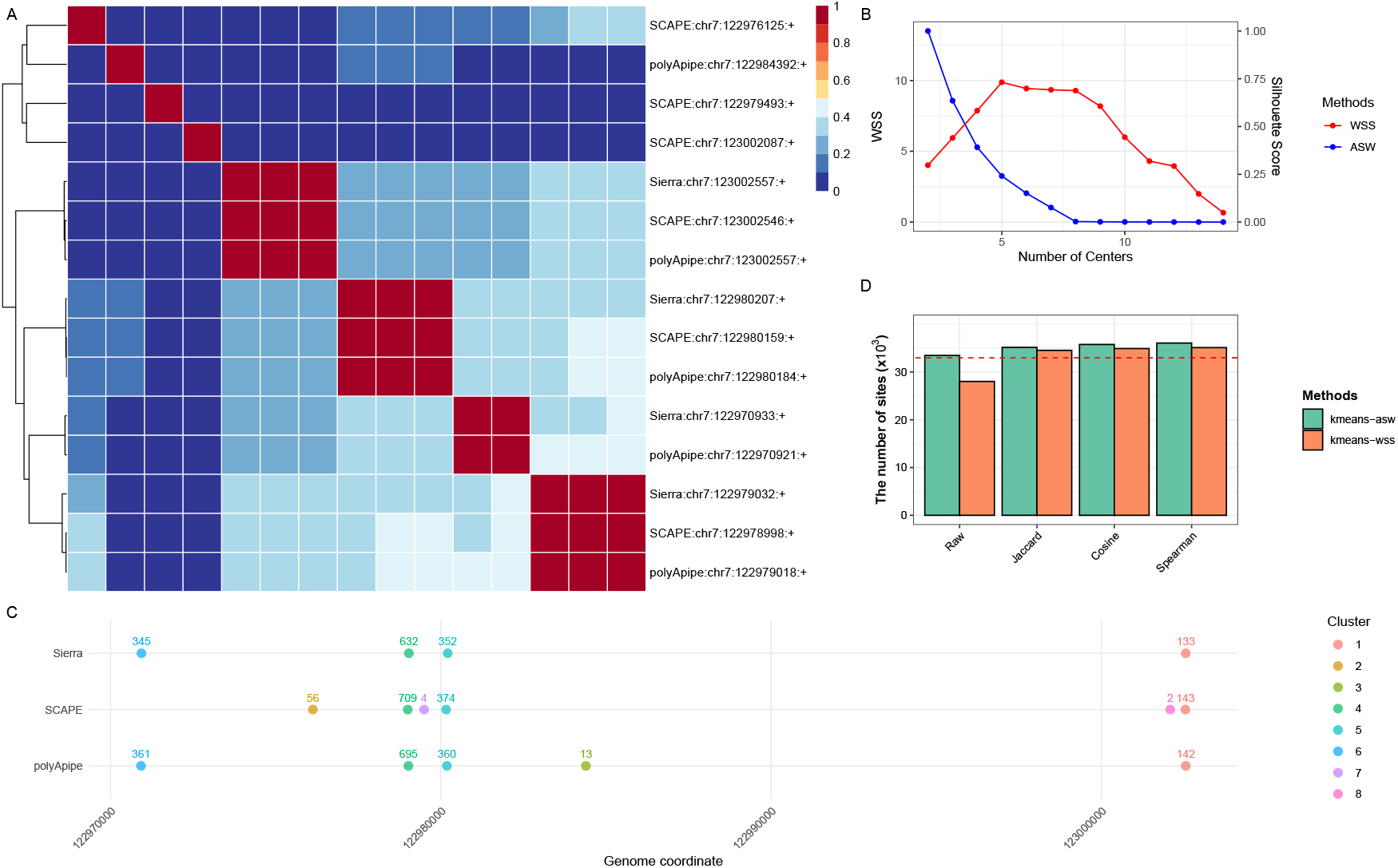
Performance of similarity-based method. A. Heatmap of cosine distance for Rbbp6 gene. B. WSS and ASW were used to decide the optimal center. C. K-means cluster results of Rbbp6 gene. D. Comparsion of different distances and methods. The red dashed line indicates the high-confidence site threshold.

Similarity-based integration, as a site alignment mechanism, is promising to mitigate the impact of site positional deviations. However, the main challenge of this strategy lies in calculating the similarity of sparse data or clustering sparse data. The site-level expression matrix is inherently sparser than the gene-level expression matrix. To assess whether different similarity metrics influence clustering outcomes, we evaluated several distance measures (Fig. 3D). Meanwhile, we evaluated whether two commonly used methods for determining the optimal number of clusters in k-means clustering, WSS and the average silhouette width (ASW), would influence clustering performance (Fig. 3D). The ASW-based method consistently yielded more high-confidence sites than the WSS-based method in all distance metrics evaluated. The largest differences between the two methods were observed when using raw counts and Euclidean distance, while the discrepancies were smaller for the Jaccard distance, Cosine distance, and Spearman correlation. All ASW-based tests exceeded the high-confidence threshold, whereas the WSS-based tests failed to reach this threshold when applied with raw counts or Euclidean distance. This may be attributed to the fact that, when using raw counts or Euclidean distance, the scale of the data and the presence of outliers can compromise the stability of the WSS metric, making the elbow point less distinguishable. In contrast, when more robust similarity measures such as cosine distance or Spearman correlation are applied, the performance of the WSS-based strategy improves significantly. Among the different distance metrics, clustering performance was generally poorer with raw counts or Euclidean distance, and more favorable when using Cosine distance or Spearman correlation.

### Similarity-based and position-based methods give comparable results

We proposed two integration strategies based on different principles. In terms of the number of high-confidence sites, both optimized strategies identified a number of sites exceeding the minimum threshold (Fig. 4A), with the highest value for the position-based strategy (160nt) slightly higher than that for the similarity-based strategy.

**Fig. 4.**
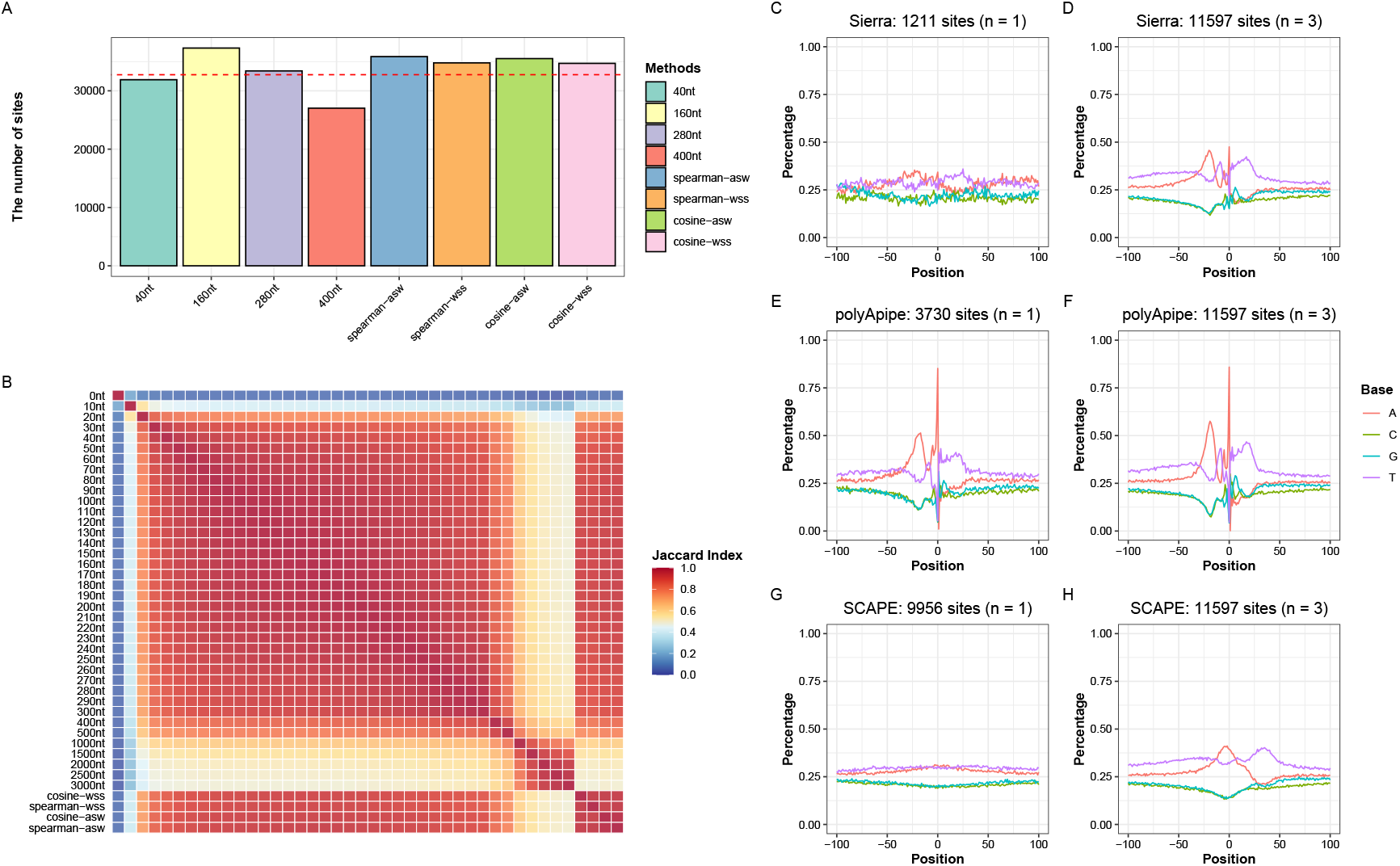
Performance comparison between position-based strategy and similarity-based strategy. A. The number of high-confidence sites identified by different methods. The red dashed line indicates the high-confidence site threshold. B. The heatmap of Jaccard index across these methods. C-H. Nucleotide composition across different methods under different confidence.

To examine whether the similarity-based strategy can alleviate some of the limitations of the position-based strategy for certain genes, we compared high-confidence sites identified by both approaches. We found that in approximately 200 genes, the similarity-based strategy partially addressed the shortcomings of the position-based strategy. Specifically, it successfully separated sites from different origins that were incorrectly merged together due to their close proximity (SFigure2), and conversely, it merged sites from the same origin that were mistakenly split apart because their distances exceeded the predefined threshold (SFigure3).

Since both strategies integrate the same data, it is expected that there should be a high proportion of overlapping sites between the acceptable results obtained by each strategy. Therefore, we used the Jaccard index to measure the overlap between different strategies (Fig. 4B). The overlap within each strategy (with different parameters) was the highest. However, the position-based strategy showed a significant reduction in the number of sites when the distance was too small (< 30nt) or too large (> 300nt), leading to a lower Jaccard index. The reduction in the number of sites stabilized between 1000 and 3000nt, so although the Jaccard index with other distances decreased, it remained relatively high with each other in this range. The similarity-based strategy showed high overlap not only between each other but also with the results of the position-based strategy within the 40-290nt range, indicating that both strategies produced comparable integration results, further validating the reliability of the integrated results. This also suggests that when employing either of the two strategies, users may not need to fine-tune parameters extensively, given the strong consistency of the main results within a certain range or under a stable distance metric.

To further validate the rationality of high-confidence sites as a measurement metric, we extracted the sequences 100bp upstream and downstream of the predicted sites from different tools and calculated the nucleotide composition at each position. If high-confidence sites reflect true signals then they are more likely to enrich the sequence characteristics of motif regions around polyadenylation sites, such as PAS, the cleavage site, the CSTF and the CF I recognition motifs.

Due to differences in site position predictions by various algorithms, even sites within the same high-confidence cluster may exhibit obvious positional deviations. Therefore, we compared the nucleotide composition differences between method-specific sites (n=1) and high-confidence sites (n=3) identified by each method (Fig 4C-H). As expected, high-confidence sites showed distinct sequence characteristics across all three methods, whereas only polyApipe method-specific sites exhibited noticeable sequence features (Fig 4E), with method-specific sites obtained by the other two methods almost completely losing these characteristics (Fig 4C and G). Theoretically, polyApipe, which relies on soft-clip reads, identifies sites with more accurate positioning and sequence features. Consequently, even method-specific sites can display prominent sequence characteristics. However, as shown in Fig 1C, polyApipe may miss a certain number of genes and sites. For high-confidence clusters, using polyApipe to characterize true sites is a good choice. For sites missed by polyApipe, if they are supported by Sierra and SCAPE, it suggests that these sites are more likely to represent true signals compared to method-specific sites. In addition, for method-specific sites, although there may be insufficient direct evidence to confirm their biological validity, supplementary information can aid in their evaluation. For instance, if a site has at least one upstream PAS and lacks features indicative of internal priming, such as an A-stretch (e.g., a sequence of seven or more consecutive adenines), it may be considered more reliable. Furthermore, if high-confidence polyA site annotations are available from the same sample, such as those derived from PAPERCLIP or direct RNA sequencing and the predicted site overlaps with an annotated site within a narrow window, this can serve as additional supporting evidence for the authenticity of the site.

### Evaluating the Extensibility of APA Integration Strategies

The similarity-based strategy offers greater extensibility, allowing the use of a wider range of similarity metrics and clustering algorithms. In Fig 5A, we demonstrate the performance of additional clustering methods, including hierarchical clustering, k-medoids, and HDBSCAN, across various similarity measures. Among these, hierarchical clustering combined with cosine distance achieved the best overall performance.

**Fig. 5.**
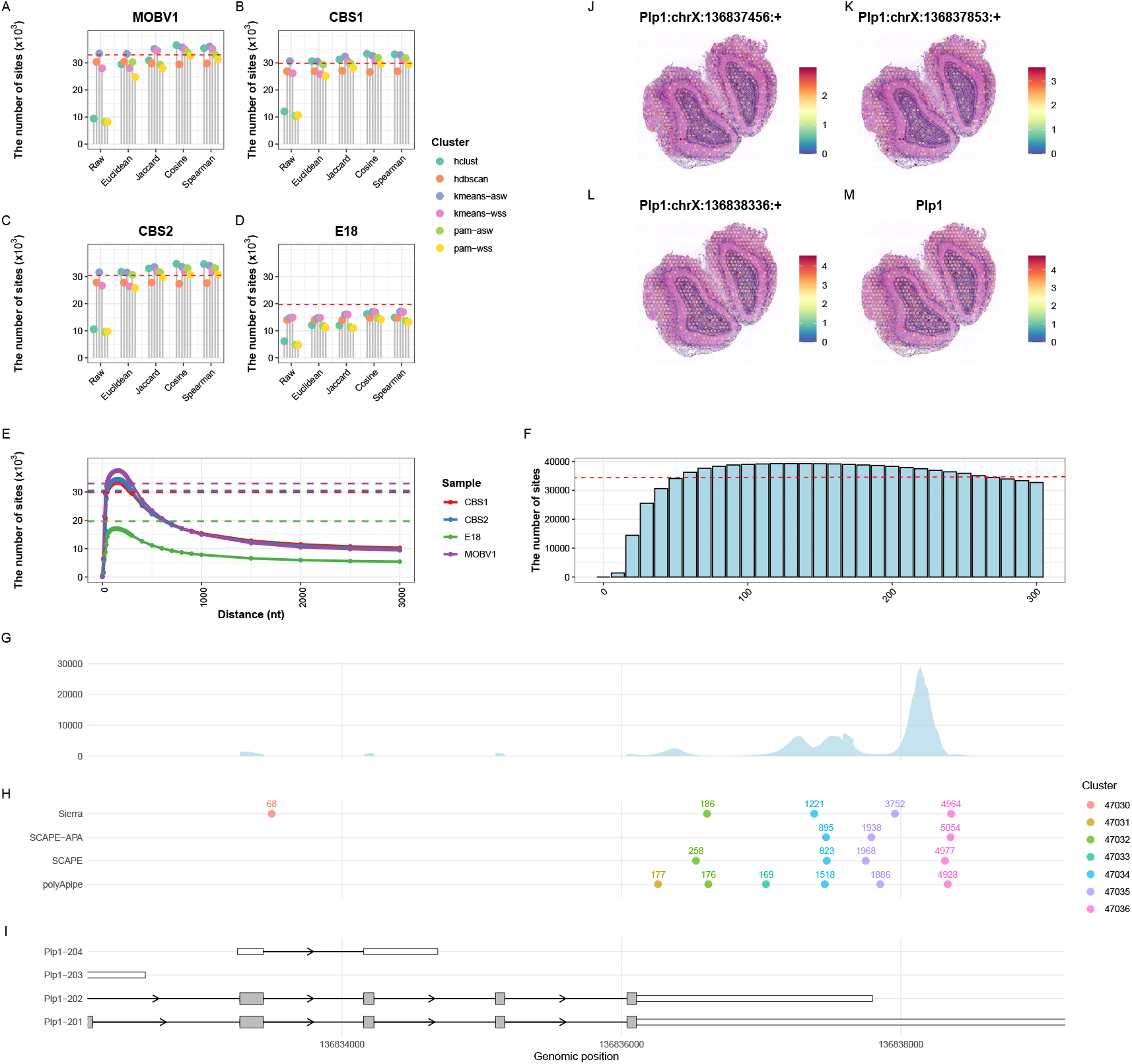
Scalability of APA Integration Strategies. A-D. The number of high-confidence sites identified by different clustering methods in MOBV1, CBS1, CBS2, E18 samples. E. The number of high-confidence sites identified by position-based method in MOBV1, CBS1, CBS2, E18 samples. F. The number of high-confidence sites obtained using the position-based integration strategy after incorporating the new method SCAPE-APA. The red dashed line indicates the high-confidence site threshold. G. Coverage profile of the Plp1 gene in the MOBV1 sample. H. Integration results from four methods. I. Genome annotation of Plp1 gene. J-M. Spatial expression patterns of the three high-confidence polyA sites in Plp1 and their correspondence with gene-level spatial expression.

We further applied both integration strategies to more samples, including CBS1, CBS2, and E18 (Fig 5B-E). For the MOBV1, CBS1, and CBS2 samples, both similarity-based and position-based strategies performed effectively. However, in the E18 sample, neither strategy surpassed the predefined threshold (high-confidence sites threshold are calculated here as the number of genes multiplied by three), likely due to the unique characteristics of this dataset, such as atypical read lengths and sequencing depth, which may have impacted the detection of the APA site, as reported in previous studies [2, 14]. When using the position-based integration strategy, we observed that the optimal distance thresholds were highly similar across all samples, showing little sample-specific variation, even in the case of the E18 sample. One possible explanation is that the effectiveness of this strategy may depend more on the current genome annotation than on the characteristics of individual samples. For instance, the median annotated 3’UTR length is 194 nucleotides. When the clustering threshold approaches or exceeds this distance, inferred polyA sites from different isoforms are more likely to be grouped into the same cluster. This observation suggests that, when applying distance-based strategies, the median 3’UTR length could serve as a biologically informed reference for selecting an appropriate clustering threshold.

In addition to the three tools (Sierra, SCAPE, and polyApipe) we initially used, our integration strategies are designed to be extensible, allowing the incorporation of additional tools. In Fig 5F, we demonstrate this by adding a new tool, SCAPE-APA, to the distance-based integration framework. SCAPE-APA is an updated version of SCAPE [16]; although it shares a similar core algorithm, improvements in read processing and the handling of splicing-induced false positives result in performance differences between the two versions. When applying all four tools to the MOBV1 sample, the integration still yielded results exceeding the high-confidence site threshold (calculated here as the number of genes multiplied by four, corresponding to the four tools used). Although integration strategies can be extended to incorporate more tools, using too many tools, especially those based on similar algorithmic principles, may not necessarily improve integration performance and could introduce redundancy.

In Fig 5G-I, we present the coverage plot of the Plp1 gene in the MOBV1 sample, along with the integration results from the four tools and the corresponding gene annotation. Plp1 gene has four annotated transcript isoforms, with distinguishable 3’UTRs. As shown in Fig 5H, the most distal site (Plp1:chrX:136838336:+) exhibits the highest expression level and overlaps exclusively with the 3’UTR of Plp1-201, suggesting that it likely originates from this isoform. In contrast, the sites Plp1:chrX:136837456:+ and Plp1:chrX:136837853:+ are located closer to the 3’UTR terminus of Plp1-202. Previous studies using ONT and in situ sequencing (ISS) technologies have demonstrated spatially distinct expression patterns of Plp1-201 and Plp1-202 in MOB (mouse olfactory bulb) [15]. Specifically, Plp1-201 is expressed in both the olfactory nerve layer (peripheral region) and the granule cell layer (central region), with particularly strong expression in the granule cell layer. In contrast, Plp1-202 is predominantly expressed in the olfactory nerve layer, with only sparse signals detected in the granule cell layer.

We examined the spatial expression patterns of the three high-confidence polyA sites. As expected, the distal site (Plp1:chrX:136838336:+) showed an expression pattern consistent with Plp1-201, being expressed in both peripheral and central regions, with stronger signals in the central region (Fig 5L). This aligns with the gene-level expression pattern (Fig 5M) and may reflect the fact that Plp1-201 is the most abundantly expressed isoform. The other two sites exhibited distinct spatial patterns more consistent with Plp1-202, with expression primarily localized to the peripheral region and markedly reduced expression in the central region, particularly for Plp1:chrX:136837853:+ (Fig 5J and K).

## Conclusion

We have developed a framework for integrating polyA sites inferred by different APA analysis tools. This framework includes two integration strategies based on different principles. The first strategy is based on genomic coordinates, clustering sites within a given distance threshold to achieve integration. The second strategy is based on expression similarity, clustering sites with similar expression patterns. By evaluating the number of high-confidence sites, their overlap sites, and sequence characteristics, we demonstrated that both strategies effectively achieve integration. Depending on the biologist’s needs, sites with varying confidence levels can be retained from the integrated results for downstream analyses, such as identifying differential sites or clustering. This integration leverages the strengths of tools with different algorithms, compensating for their weaknesses, and provides a more comprehensive APA landscape.

The position-based strategy is straightforward and simple, with the main challenge being the selection of an appropriate distance. As the distance increases, the number of high-confidence sites initially increases and then decreases. A distance that is too small may fail to cluster sites that should be grouped together, while a distance that is too large may result in too many sites being clustered together. Even at a distance that maximizes the number of high-confidence sites, there is still a risk that sites within the same cluster may originate from different signals, but are incorrectly grouped. Fortunately, the results perform well across a broad range (50–290 nt in mouse), with particularly high consistency. This indicates that users may not need to engage in extensive fine-tuning of the distance threshold. Besides, based on our results, we do not recommend setting the distance parameter beyond the median length of the 3’UTR for the species.

The similarity-based strategy is novel, and we designed a new framework allowing users to calculate different similarities and use various clustering methods according to their needs. The main challenge of this strategy is clustering sparse data. We employed different distance metrics, such as cosine distance, Jaccard distance, and Spearman correlation coefficient, and used the distance matrix for clustering. This approach produced superior results when employing stable distance metrics, as compared to using raw counts or Euclidean distance. The choice of clustering method and the method for determining the number of clusters are also worth exploring. Some methods tend to produce larger clusters, which are not suitable for our case. While ASW-based center determination typically exhibits superior performance compared to WSS, both approaches produce robust and highly consistent outcomes under stable distance metrics. As a result, users may not need to invest significant effort in parameter adjustment.

Furthermore, we showcased the extensibility of the integration framework, which accommodates a wider array of APA tools, supports diverse clustering strategies and distance measures, and is readily applicable to other datasets. Additionally, analysis of the spatial expression profile of the Plp1 gene revealed that the high-confidence sites identified by our method display coherent spatial patterns, further underscoring the capacity of our integrative framework to uncover biologically relevant information.

However, three challenges require additional attention. First, although high-confidence sites theoretically reflect true signals, these signals may originate from genuine transcripts or artificially shortened transcripts, i.e. bias shared by all the methods. Therefore, additional tools may be needed in the future to remove common bias for biological studies. Second, we observed that many genes are still missed by all methods. For example, 1563 genes were identified only by Space Ranger, and 1420 genes were identified by both Space Ranger and ONT, but missed by the other three methods. This suggests that there is still room for new APA identification methods. Even though many of these genes may be lowly expressed, mitochondrial in origin, or derived from non-coding RNAs, they may still carry valuable information that warrants further investigation. Third, while high-confidence sites help eliminate false positives, addressing false negatives remains challenging and currently relies on additional sources of information, such as complementary datasets or specific sequence features, highlighting the need for a unified and comprehensive approach.

## Competing interests

No competing interest is declared.

## Author contributions statement

MR and QZ conceptualized the study. QZ performed data analysis and wrote the manuscript. MR reviewed the manuscript.

## Funding

MR was supported by a Wellcome Trust Discovery Award (Ref 227415/Z/23/Z). QZ was supported by a University of Manchester-China Scholarship Council Joint Scholarship.

## Acknowledgments

We thank the anonymous reviewers for their valuable suggestions. We thank the authors of the datasets used in this study. We also appreciate the constructive discussions with all members of the Rattray Lab.

## Notes

### Competing Interest Statement

The authors have declared no competing interest.

