## Supplementary figures and images for "metaAPA: a tool for integration of PolyA site predictions from single-cell and spatial transcriptomics"

### Sfigure1_other_sample_upset_plots_v1.pdf

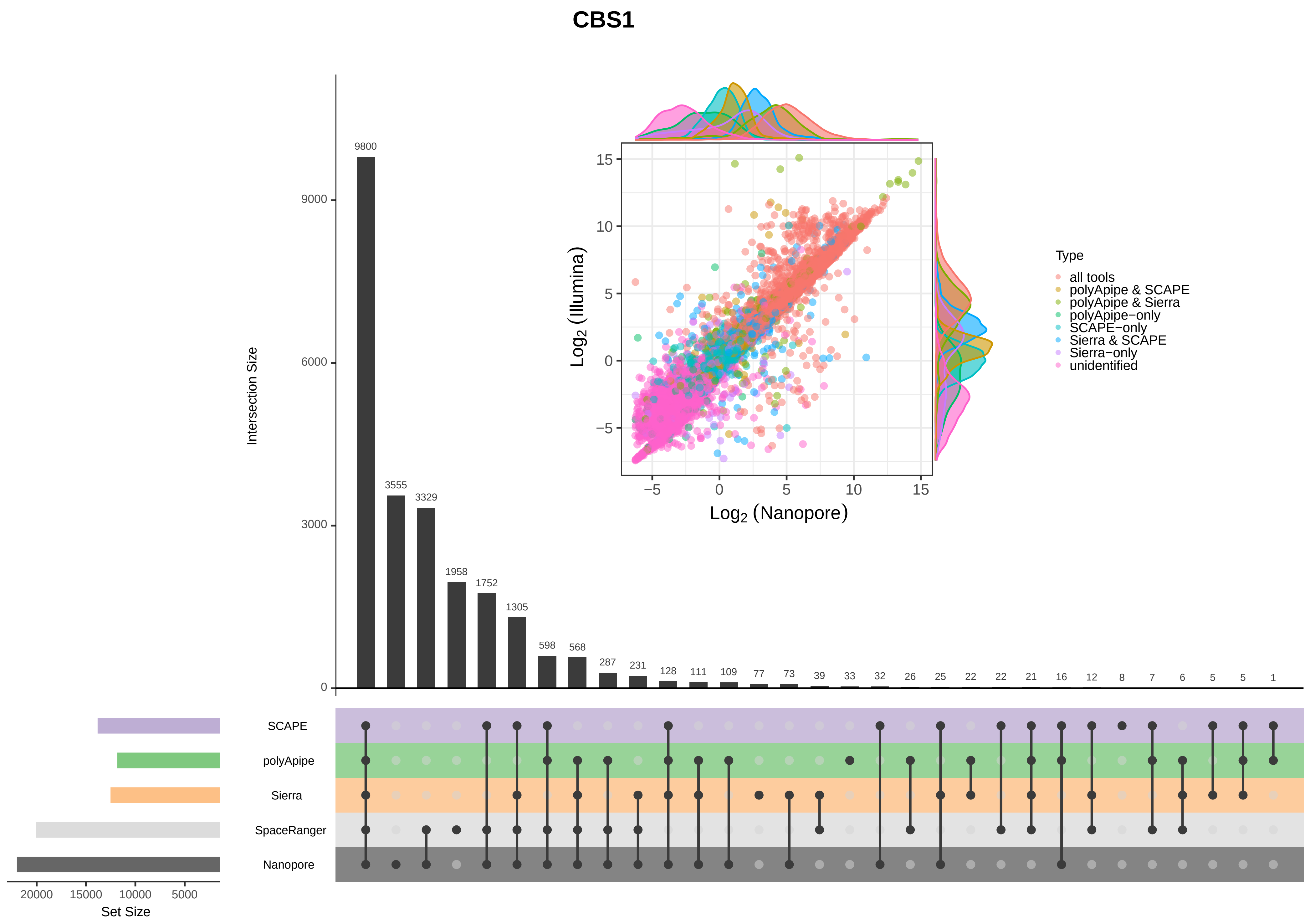

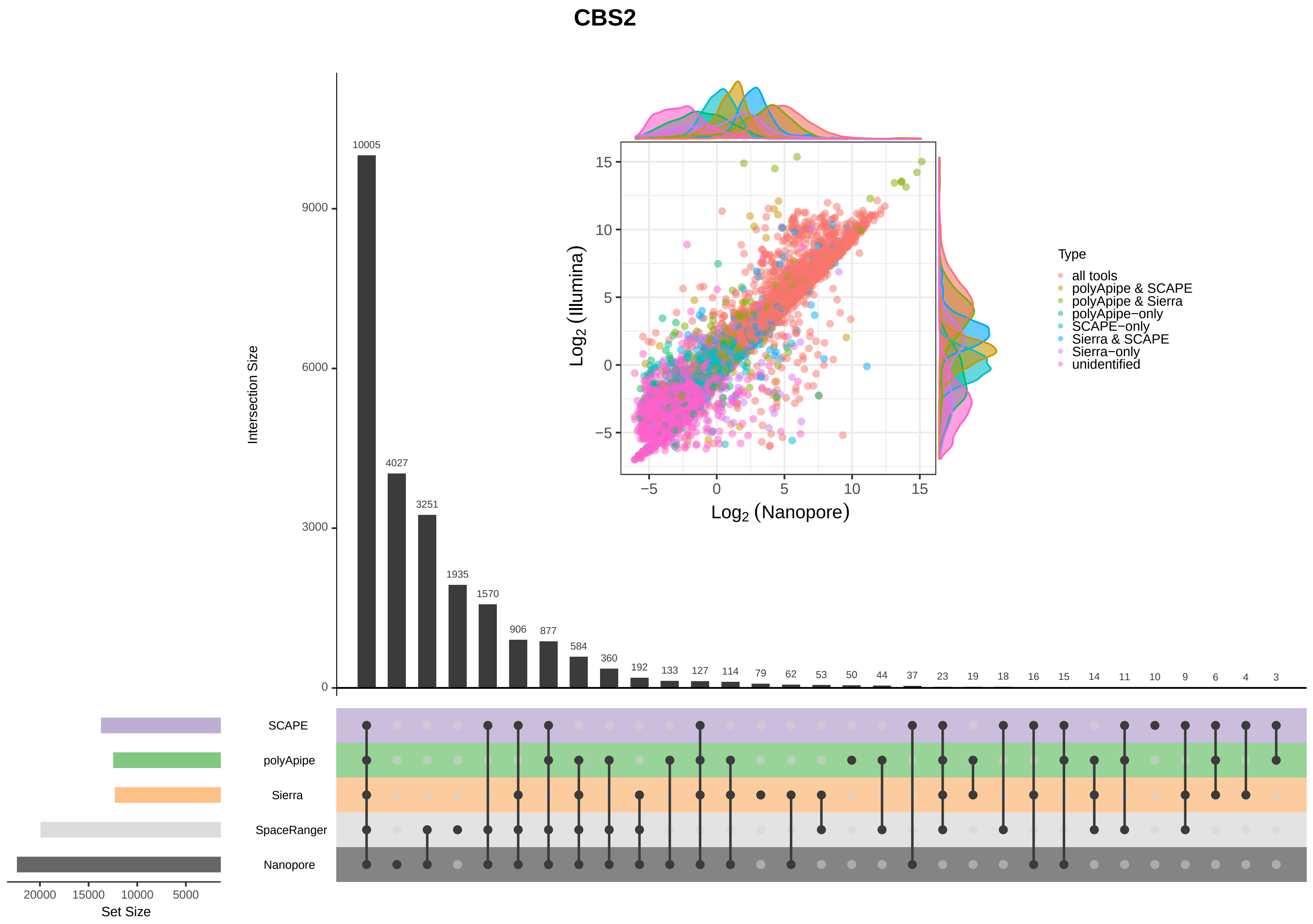

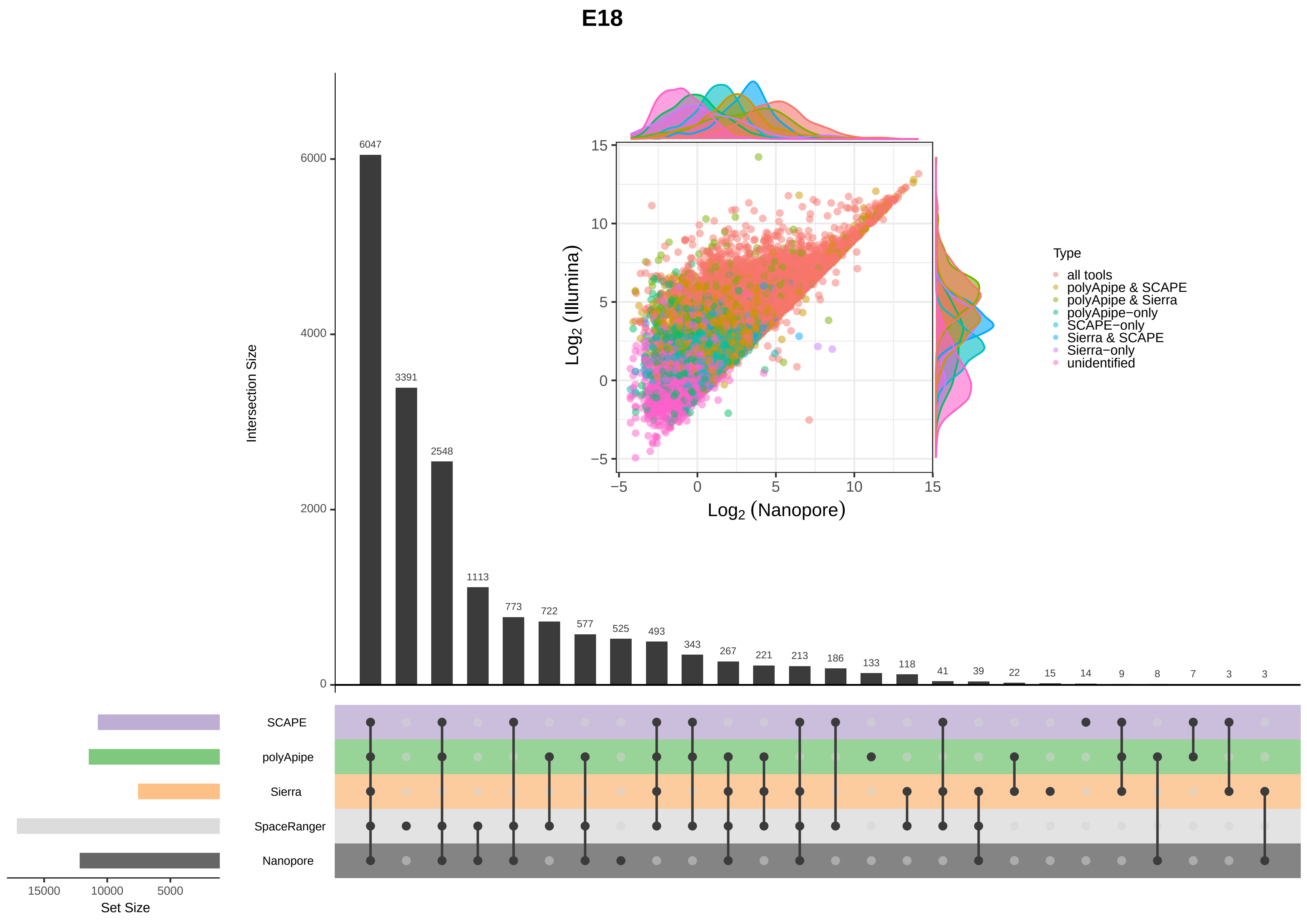

### Sfigure2_MOBV1_Rbis_coverage_alignment_plot.pdf

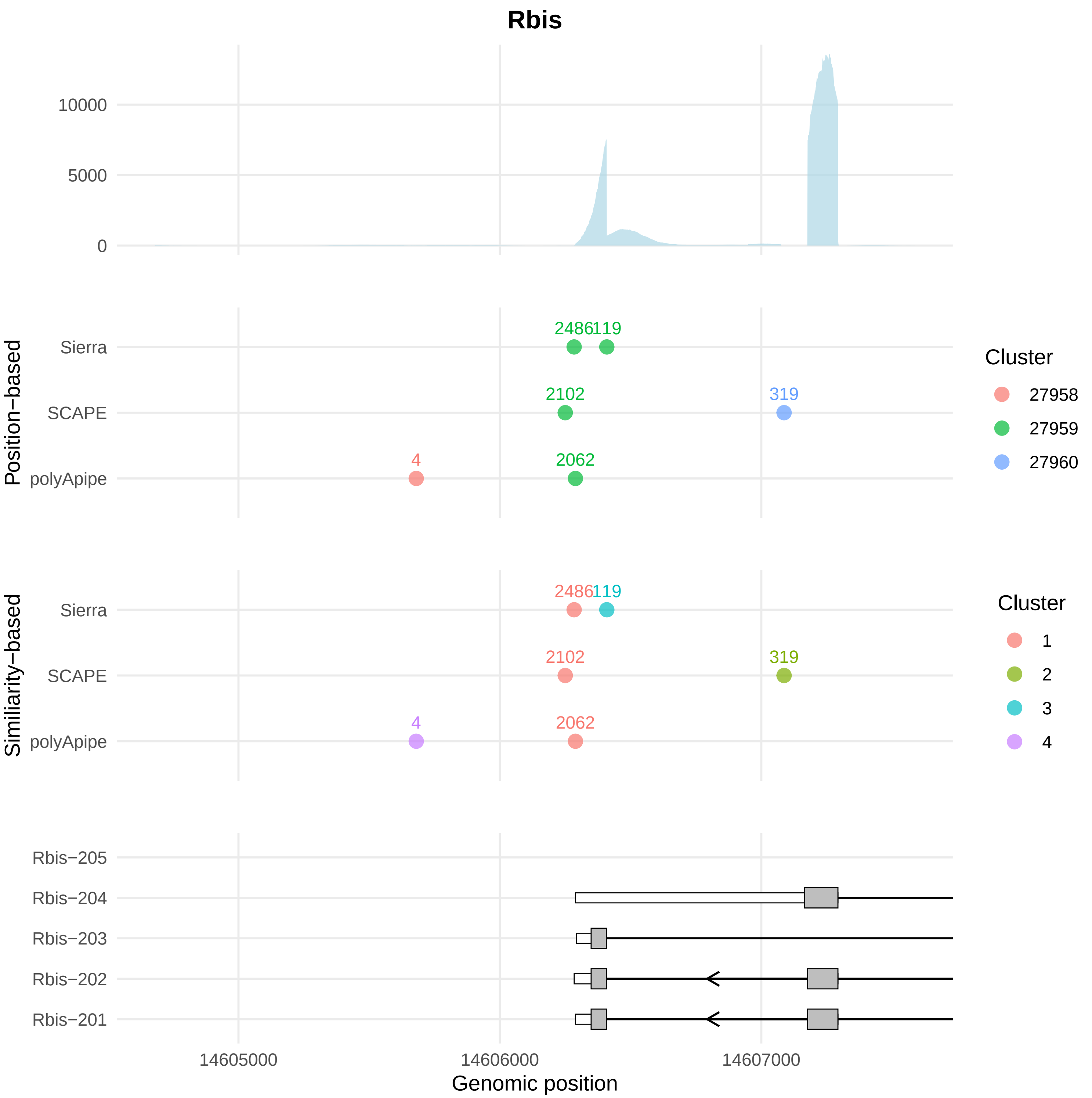

### Sfigure3_MOBV1_Sox8_coverage_alignment_plot.pdf

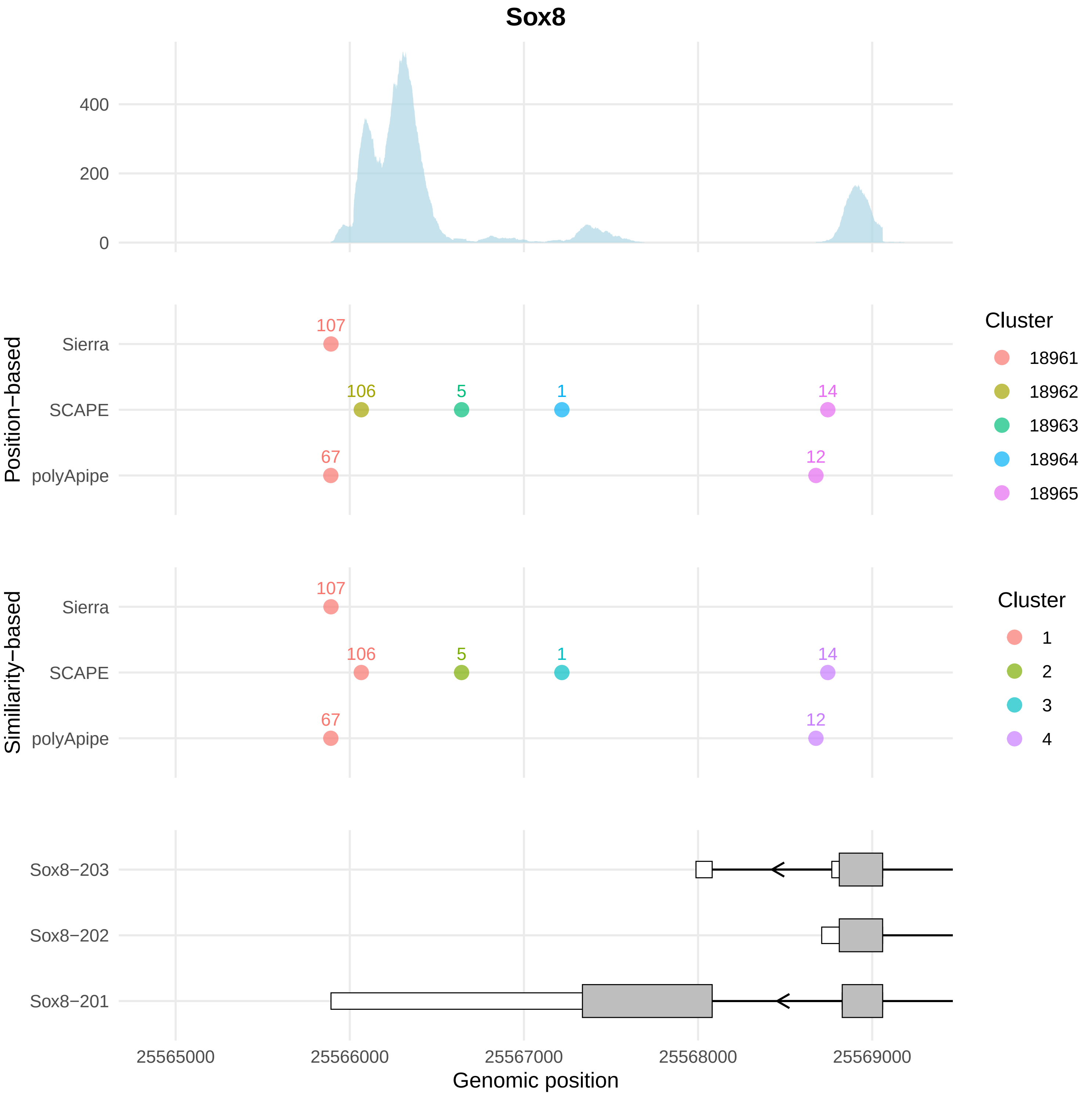
